# Combining a rhesus cytomegalovirus/SIV vaccine with neutralizing antibody to protect against SIV challenge in rhesus macaques

**DOI:** 10.1101/2025.03.20.644395

**Authors:** Jessica Coppola, Mara Parren, Raiza Bastidas, Karen Saye, Jacqueline Malvin, Joseph G. Jardine, Roxanne M. Gilbride, Sohita Ojha, Shana Feltham, David Morrow, Aaron Barber-Axthelm, Rachele Bochart, Randy Fast, Kelli Oswald, Rebecca Shoemaker, JeJrey D. Lifson, Louis J. Picker, Dennis R. Burton, Scott G. Hansen

## Abstract

A vaccine is widely regarded as necessary for the control of the HIV pandemic and eventual eradication of AIDS. Neutralizing antibodies and MHC-E-restricted CD8+ T cells have both been shown capable of vaccine protection against the simian counterpart of HIV, SIV, in rhesus macaques. Here we provide preliminary evidence that combining these orthogonal antiviral mechanisms can provide increased protection against SIV challenge such that replication arrest observed following vaccination with a rhesus cytomegalovirus (RhCMV/SIV)-based vaccine was enhanced in the presence of passively administered incompletely protective levels of neutralizing antibody. The report invites studies involving larger cohorts of macaques and alternate routes of providing neutralizing antibody.

## Introduction

Only two vaccination modalities have been shown to consistently provide protection against human immunodeficiency virus (HIV) and/or its simian counterpart, simian immunodeficiency virus (SIV). The first is neutralizing antibodies. Autologous neutralizing antibodies induced through vaccination with a stabilized recombinant envelope (Env) molecule have been shown to provide protection against challenge with the corresponding HIV/SIV chimera (SHIV) in rhesus macaques (RMs) (Pauthner et al., 2017; Petitdemange et al., 2019). In addition, passively administered broadly neutralizing antibodies have been shown to provide protection against SHIV and SIV in RMs (Parren et al., 2001; Pegu et al., 2019; Zhao et al., 2022) and against HIV in humans (Corey et al., 2021). The second modality is the MHC-E-restricted CD8+ T cell-targeted rhesus cytomegalovirus (RhCMV/SIV)-based vaccine. When properly genetically programmed (strain 68-1 and certain derivatives), RhCMV/SIV vaccine vectors elicit MHC-E-restricted CD8+ T cells in RMs and these responses mediate early complete arrest of SIVmac239 replication in about 60% of RM after repeated, limiting dose SIV challenge and the vast majority of these protected RMs will subsequently clear SIV infection completely (Hansen et al., 2011; Hansen et al., 2013a; Hansen et al., 2013b; Hansen et al., 2016; Hansen et al., 2019; Malouli et al., 2021; Verweij et al., 2021; Picker et al., 2023). RhCMV/SIV vaccines (typically containing Gag, Rev/Tat/Nef, and 5’-Pol inserts) do not induce antibodies and are effective in the absence of Env expression (Hansen et al., 2011; Hansen et al., 2013a; Hansen et al., 2019; Malouli et al., 2021; Hansen et al., 2022) They therefore represent a mode of protection completely orthogonal relative to nAbs (Hansen et al., 2011; Hansen et al., 2013a; Hansen et al., 2019; Malouli et al., 2021; Hansen et al., 2022). The MHC-E-restricted CD8+ T cell responses are very durable, with protective efficacy shown up to 10 years post-primary vaccination in RMs (Picker et al., 2023).

An important distinction exists between the nature of protection mediated by nAbs and the MHC-E-restricted CD8+ T cell responses described as above. For nAbs, protection is generally observed as an early sterilizing type of immunity, with very little or no viral replication or spread observed following challenge (Hessell et al., 2007; Hessell et al., 2010; Barouch et al., 2013; Burton, 2023; Stab et al., 2023). For RhCMV/SIV-induced MHC-E restricted CD8+ T cell-mediated protection in RMs, some SIV replication and spread occurs early after effective challenge as evidenced by direct viral measurements in tissues in the first few weeks following and by the post-challenge induction of T cell responses to SIV antigens (Ags) not included in the vaccine (e.g., anti-Vif and anti-Env T cell responses) (Hansen et al., 2013a; Picker et al., 2023). Infection may also result in “blips” in plasma virus and can be demonstrated by adoptive transfer of hematolymphoid cells to naive RMs resulting typical SIV infection (Hansen et al., 2013a; Hansen et al., 2019). In RhCMV/SIV-vaccine-protected RMs, this early take of infection is stringently controlled (“replication arrest”) and SIV eventually is cleared (Hansen et al., 2011; Hansen et al., 2013a; Hansen et al., 2019; Picker et al., 2023).

Both of these effective protective modalities have important limitations. The serum nAb titers required for sterilizing immunity against (S)HIV (and indeed against many viruses) are high, typically in the many hundreds (Pauthner et al., 2019; Pegu et al., 2019; Saunders et al., 2022; Zhao et al., 2022) and such titers are challenging to induce and sustain by vaccination. Although the RhCMV/SIV vaccine is very durable it still only protects ∼60% of SIV-challenged RMs (Hansen et al., 2011; Hansen et al., 2013a; Hansen et al., 2019; Picker et al., 2023). A key question is then whether these two modalities may synergize, in particular when nAb titers wane below the level required to completely stop take of infection. Specifically, does the presence of nAbs at sub-completely protective (sub-threshold) neutralizing titers at the time of challenge increase the proportion of RhCMV/SIV vaccinated RMs undergoing replication arrest? Put alternatively, can lower nAb titers contribute to protection in the presence of an MHC-E-restricted T cell response to HIV?

We investigated these questions in a pilot study in the RM model using a 68-1 RhCMV/SIV vaccine, passively administered anti-SIV nAb and SIVmac239 challenge. Passive antibody was chosen since no SIV Env targeted vaccine has yet been developed that can reliably induce nAbs against SIVmac239. We first determined the serum titer of nAb alone that would provide complete protection against SIVmac239 challenge since we wished to use a sub-protective titer in the synergy experiment. We then vaccinated two cohorts of RMs with the RhCMV/SIV vaccine and 73 weeks later administered nAb or control Ab followed by challenge with high-dose SIVmac239. We used a high-dose SIV challenge because of the logistics of the experiment to keep the number of passive antibody administrations manageable. The results from this pilot study are promising, indicating potential synergy between nAbs and MHC-E-restricted CD8+ T cells in resisting SIV infection, even under conditions of stringent high dose viral challenge.

## Materials and Methods

### Rhesus macaques

30 Indian RMs (Macaca mulatta) were used in this study. All RMs were classified as specific pathogen free defined by being free of cercopithecine herpesvirus 1, D-type simian retrovirus, simian T-lymphotropic virus type 1, SIV, and M. tuberculosis but naturally infected with RhCMV at study initiation. All RMs used in this study were housed at the Oregon National Primate Research Center (ONPRC) in Animal Biosafety Level 2+ rooms. RM care and all experimental protocols and procedures were approved by the ONPRC Institutional Animal Care and Use Committee (IACUC). The ONPRC is a category I facility. The Laboratory Animal Care and Use Program at the ONPRC is fully accredited by the American Association for Accreditation of Laboratory Animal Care and has an approved assurance (no. A3304-01) for the care and use of animals on file with the National Institutes of Health Office for Protection from Research Risks. The IACUC adheres to national guidelines established in the Animal Welfare Act (7 U.S.C. Sections 2131–2159) and the Guide for the Care and Use of Laboratory Animals (eighth edition) as mandated by the US Public Health Service Policy. RMs were separated as follows: Group 1 (68-1 RhCMV/SIV vaccinated and K11-LS infused RMs, *n* = 9), Group 2 (CMV/SIV vaccinated and DEN3 infused RMs, *n* = 9), Group 3 (K11-LS infused RMs, *n* = 6). For this study both DEN3 and K11-LS were administered intravenously (IV) at a controlled rate by trained personnel in accordance with the study IACUC approved protocol, with no adverse events observed.

### Cell lines

TZM-bl cells (NIH AIDS Reagent Program) were used in a pseudovirus neutralization assay. Human HEK 293 T cells (ATCC) were used for pseudovirus production. Expi293F cells (ThermoFisher) were used for monoclonal antibody production.

### SIV pseudovirus production

SIV pseudovirus Env construct was co-transfected with Env-deficient backbone plasmid (pSG3ΔEnv) in a 1:2 ratio with transfection reagent FuGENE 6 (Promega) in HEK 293 T cells according to the manufacturer’s instructions. After 72 hours of transfection, supernatants containing viruses were harvested and sterile filtered (0.22 μm) (EMD Millipore) and frozen at −80°C for long-term storage.

### Antibody production and characterization

Antibody heavy chain (HC) and light chain (LC) constructs were transiently expressed with the Expi293 Expression System (ThermoFisher). HC and LC plasmids were cotransfected at a 1:2.5 ratio with transfection reagent FectoPRO (Polyplus) in Expi293 cells according to the manufacturer’s instructions. After 24h, cells were fed with 300 mM valproic acid and 40% glucose (Gibco). After 5 days of transfection, cell supernatants were harvested and sterile filtered (0.22 μm). Antibody was purified by Protein A Sepharose (GE Healthcare) as described previously (Sok et al., 2013). Antibody batches were viewed on an analytical high-performance liquid chromatography (HPLC) system (Agilent 1260 Infinity II) to confirm correct size and purity, analyzed in neutralization assays to confirm batch-to-batch potency, and endotoxin tested prior to pooling all batches and freezing at -80°C for long-term storage or shipment to OHSU.

### TZM-bl neutralization assay

Serially diluted heat-inactivated serum or antibody was incubated with SIVmac239 pseudovirus or murine leukemia (MLV) (negative control) pseudovirus in half-area 96-well white plates using Dulbecco’s Modified Eagle Medium (DMEM) (Gibco) supplemented with 10% FBS, 2 mM L-glutamine (Gibco), 100 U/mL Penicillin/Streptomycin (Gibco). K11-LS and DEN3 were included as positive and negative antibody only controls, respectively. After 1 hour incubation at 37°C, TZM-bl cells with 20 μg/mL DEAE-dextran were added onto the plates at 10,000 cells/well. Final antibody concentrations for the dilution series were calculated based on the total volume of the assay (antibody or serum + virus + TZM-bl cells). After 72 hours of incubation, culture supernatants were removed, and cells were lysed in luciferase lysis buffer (25 mM Gly-Gly pH 7.8, 15 mM MgSO_4_, 4 mM EGTA, 1% Triton X-100). Luciferase activity was measured by adding BrightGlo (Promega) according to the manufacturer’s instructions. Assays were tested in duplicate wells and independently repeated at least twice. Neutralization IC50 or ID50 titers were calculated using “One-Site Fit LogIC50” regression in GraphPad Prism 10.

### T cell assays

ICS was performed as described below. SIV-specific CD4+ and CD8+ T cell responses were measured in PBMC by flow cytometric ICS as previously described (Hansen et al., 2011; Hansen et al., 2013a; Malouli et al., 2021; Hansen et al., 2022). T cell responses to total SIV antigens were measured using mixes of sequential 15-mer peptides (11 amino acid overlap) spanning the SIVmac239 Gag, Pol, Nef, Rev, Tat, and Vif proteins. Mononuclear cells were stimulated in the presence of antibodies anti-CD28 (CD28.2, Purified 500 ng/test; Life Tech, CUST03277) and anti-CD49d (9F10, Purified 500 ng/test; Life Tech, CUST03278) then were incubated at 37°C with individual peptides or peptide mixes and antibodies for 1h, followed by an additional 8h incubation in the presence of Brefeldin A (5 μg ml^−1^; BioLegend, 91850). Stimulation in the absence of peptides served as background control.

After incubation, stimulated cells were stored at 4°C until staining with combinations of fluorochrome-conjugated monoclonal antibodies including: anti-CD3 (SP34-2: PacBlue, BD Biosciences, 624034), anti-CD4 (L200; BV510, BD Biosciences, 624340), anti-CD8α (SK-1: Life Technologies, CUST04424), anti-CD69 (FN50: PE/Dazzle594, BioLegend, 93437), anti-IFNγ (B27: APC, BioLegend, 96019), anti-TNFα (Mab11: PE, BioLegend, 96019), anti-Ki67 (B57: BD Biosciences, 624046). For memory phenotyping from whole blood, the following antibodies were used: anti-CCR5 (3A9: APC, BD Biosciences, 624346), anti-CCR7 (G043H7: Biotin, BioLegend, 93747), Streptavidin (BUV496, BD Biosciences, 624283), anti-CD20 (2H7: APC-Fire 750, BioLegend, 93924), anti-CD28 (CD28.2: PE/Dazzle594, BioLegend, 93924), anti-CD3 (SP34-2, BUV395, BD Biosciences, 624310), anti-CD8β (2ST8.5H7: BUV563, BD Biosciences, 624284), anti-CD25 (2A3: BUV737, BD Biosciences, 624286), anti-CXCR5 (MU5UBEE: SuperBright436, Life Technologies, 62-9185-42), anti-CD95 (DX2: BV605, BioLegend, 93384), anti-CD69 (FN50: BV650, BioLegend, 93755), anti-CD8α (RPA-T8: BV711, BioLegend, 900006277), anti-PD-1 (eBioJ105: SuperBright780, Life Technologies, 78-2799-42), anti-γδTCR (B1; PerCP-eFluor710, BioLegend, 900002746), anti-CD127 (HIL-7R-M21: PE, BD Bioscience, 624048). anti-HLA-DR (L243: PE/Dazzle 594, BioLegend, 93957), anti-CD4 (L200: BV510, BD Biosciences, 624340), anti-Ki67 (B57: FITC, BD Biosciences, 624046).

Stained samples were analyzed on an LSR-II or FACSymphony A5 flow cytometer (BD Biosciences). Data analysis was performed using FlowJo software (BD Biosciences). In all analyses, gating on the lymphocyte population was followed by the separation of the CD3+ T cell subset and progressive gating on CD4+ and CD8+ T cell subsets. Antigen-responding cells in both CD4+ and CD8+ T cell populations were determined by their intracellular expression of CD69 and either or both of the cytokines IFN-γ and TNFα. Assay limit of detection was determined as previously described (Hansen et al., 2011; Hansen et al., 2013a; Hansen et al., 2019; Malouli et al., 2021; Hansen et al., 2022), with 0.05% after background subtraction being the minimum threshold used in this study. After background subtraction, the raw response frequencies above the assay limit of detection were “memory-corrected” (e.g., % responding out of the memory population), as described (Hansen et al., 2011; Hansen et al., 2013a; Hansen et al., 2019; Malouli et al., 2021; Hansen et al., 2022). For memory phenotype analysis, CD4+ or CD8+ T cells were subdivided into the memory subsets of interest on the basis of surface phenotype (CD28 vs. CD95), with memory defined as CD28+/- and CD95+.

### SIVmac239 challenge experiments

To define K11-LS half-life, first 2 RMS were treated with 20 mg/kg, neutralizing titers allowed to decay, administered a second dose of 10 mg/kg and then half-life determined. A second group of 4 RMs was treated with 20 mg/kg K11-LS and then 10mg/kg of K11-LS (Figure 1). All RMs were then challenged with a dose of 900 focus-forming units (FFU) of SIVmac239.

**Figure 1.**
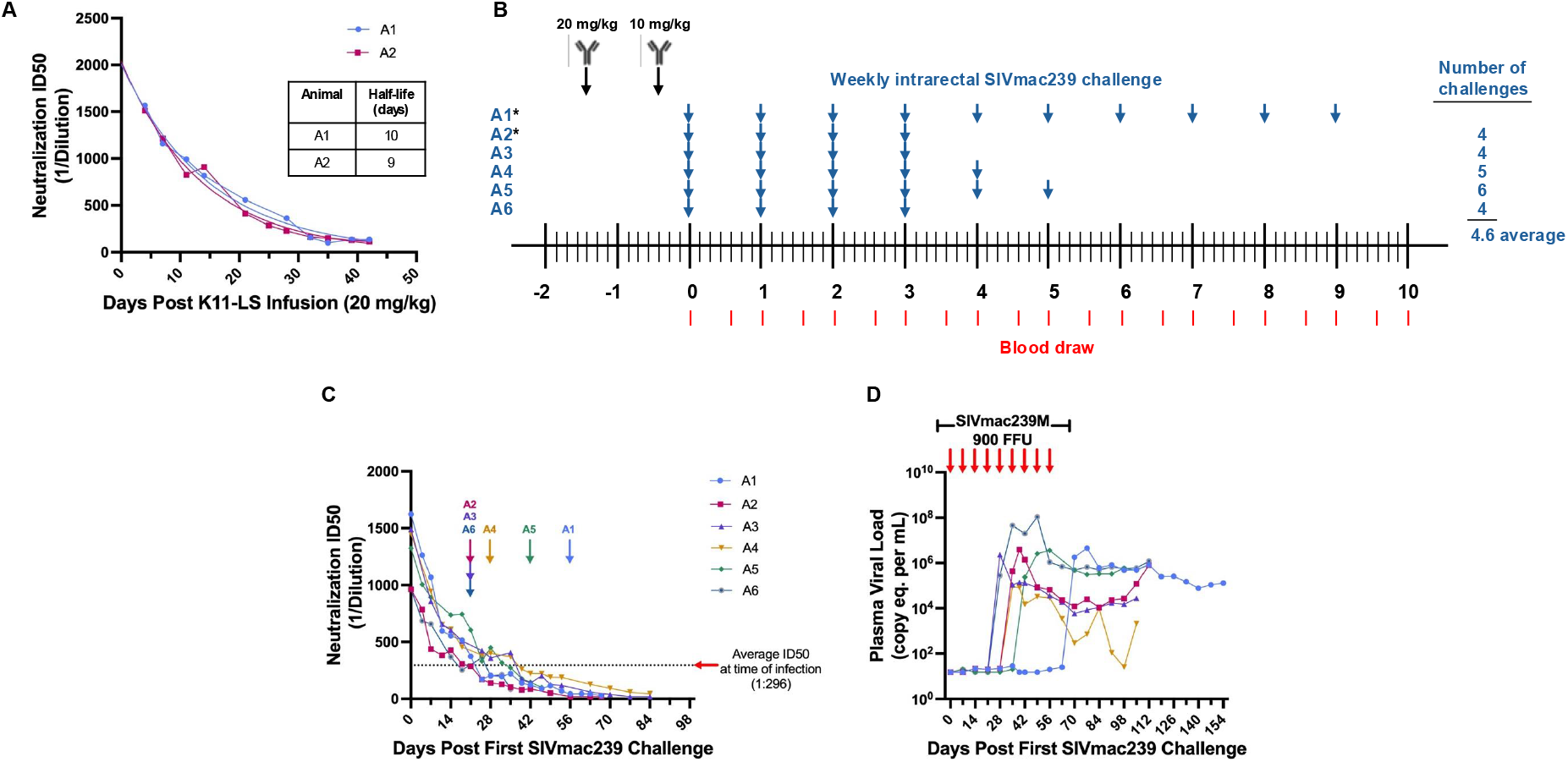
A neutralization ID50 of 1:300 is the threshold for protection against high-dose (900 FFU) SIVmac239 challenge by antibody K11-LS. (A) The half-life of K11-LS was determined to be 9.5 days after two RMs (A1 and A2) were administered 20 mg/kg and titers were allowed to decay. (B) RMs A4-A6 were administered 20 mg/kg K11-LS on day -10 followed by 10 mg/kg on day -3. *Animals A1 and A2 were administered 20 mg/kg K11-LS and titers were allowed to decay to 1:100 or less prior to a second infusion of 10 mg/kg at day 42 to calculate half-life. All RMs were challenged 3 days after the second K11-LS infusion and every week thereafter until infected (blue arrow). Red lines indicate blood draws. (C) Neutralization ID50s against SIVmac239 PSV were on average 1:296 at time of infection, with an average of 4.6 challenges before infection. This excludes animal A1, who became infected after 9 challenges with an ID50 of 1:44, thus, exhibiting an inherent resistance to SIVmac239. Arrows indicate the last challenge before infection. (D) Viral load in all 6 RMs shown in panel C post first effective 900 FFU SIVmac239 challenge. Arrows indicate weekly 900 FFU challenge.

Animals were challenged weekly until documented take of infection in terms of sustained plasma viremia. The SIVmac239 stock was titered using the CMMT-CD4-LTR-β-Gal sMAGI cell assay (National Institutes of Health AIDS Reagent Program). The dose of 900 FFU was selected based on titering experiments where 100% of unvaccinated, untreated animals became infected after 2 challenges. For the combined K11-LS/T cell protection studies, at the end of the vaccine phase, all vaccinated and unvaccinated RMs were SIV challenged intra-rectally with a dose of 900 FFU of SIVmac239 until infection could be documented as either onset of sustained plasma viremia and/or *de novo* development of CD4+ and CD8+ T cell responses to SIV Vif), at which time challenge was discontinued, as previously described (Hansen et al., 2011; Hansen et al., 2013b; Hansen et al., 2019).

### Viral load measurement

Plasma SIV RNA levels were determined using an SIV Gag-targeted quantitative RT-PCR format assay, with 6 replicate reactions analyzed per extracted sample for an assay threshold of 15 SIV RNA copies/ml, as previously described (Bolton et al., 2016).

## Results

### Passive transfer studies of K11 to determine conditions for synergy experiment

The monoclonal nAb K11 was originally isolated from a SIVmac239-infected RM, binds to a glycan hole on gp120, and neutralizes SIVmac239 at an IC50 of 100 ng/ml (Zhao et al., 2022). This high potency makes it a valuable tool for investigating potential synergy with other immune responses. A half-life extended version (Ko et al., 2014) of K11, K11-LS, was generated and its half-life was determined to be approximately 9.5 days in RMs, as assessed from neutralization ID50s against SIVmac239 pseudovirus (PSV), following a single 20 mg/kg infusion of K11-LS (Figure 1A). In a preliminary study, RMs were administered 20 mg/kg of K11-LS, allowing titers to decay to 1:100 or lower before administering a second dose of 10 mg/kg, followed by weekly intrarectal challenge with 900 FFU SIVmac239M (Fennessey et al., 2017; Khanal et al., 2019) (Figure 1B). RMs became infected after 4.6 challenges on average, excluding animal A1, which became infected after 9 challenges (Figure 1B, C, D). The study indicated that maintaining a neutralization ID50 above 1:300 prevents SIVmac239 infection for most RMs (Figure 1C, D). Thus, to assess synergy with the RhCMV/SIV vaccine, nAb titers needed to be below 1:300 prior to SIVmac239 challenge.

It is worth noting that this protective threshold differs somewhat from the previously reported ID50 of 1:609 necessary for protection when RMs were challenged intravenously with low-dose SIVmac239 (Zhao et al., 2022). This may reflect the use of a different challenge dose and/or different challenge routes used in the two studies. To achieve a neutralization ID50 of 1:200 within 21 days, different K11-LS doses – 4 mg/kg, 3 mg/kg, and 2 mg/kg – were administered to nine RMs with neutralization titers determined twice weekly. A 3 mg/kg dose was determined to be optimal for achieving the desired ID50 within the given time frame (Figure 2).

**Figure 2.**
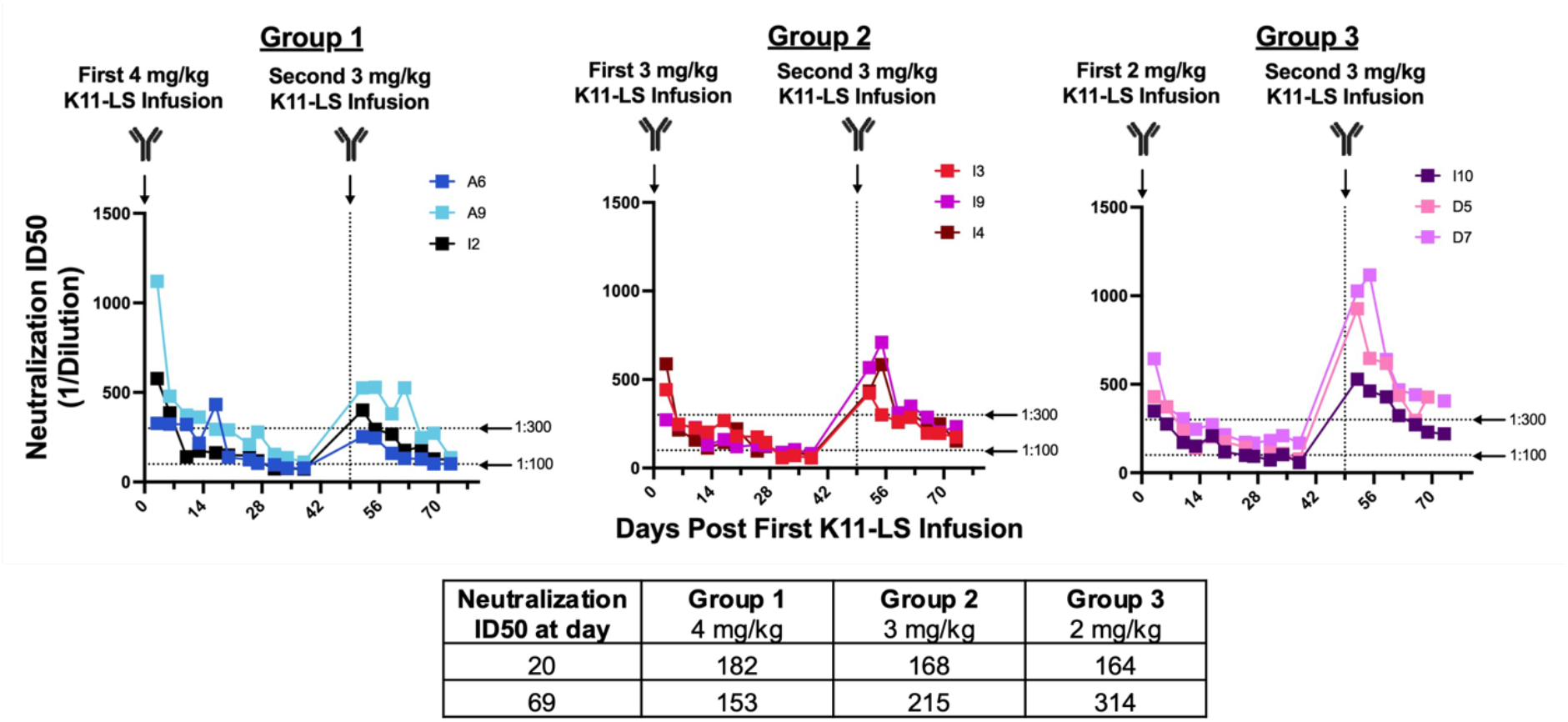
Administering K11-LS at a dose of 3 mg/kg achieves a neutralization titer between 1:100 and 1:300 within 21 days. RMs were administered 4 mg/kg (Group 1), 3 mg/kg (Group 2), or 2 mg/kg (Group 3) on day 0. 20 days later, the geometric mean ID50s were within the desired range of 1:100 – 1:200. A second dose of 3 mg/kg K11-LS was administered to all 3 groups on day 49 as the 3 mg/kg group exhibited the least variability between animals. 20 days after the second K11-LS infusion, geometric mean ID50s were 1:153, 1:215, and 1:314 for Groups 1, 2, and 3, respectively. A dose of 3 mg/kg of K11-LS was chosen for subsequent studies as ID50s were within the desired range and were the most consistent between animals.

### The frequency of replication arrest efficacy was higher with combined RhCMV/SIV vaccine and passive suboptimal neutralizing antibody than with vaccine alone

In the next study, to investigate potential ability of incompletely protective levels of nAb to combine with RhCMV/SIV vaccination for increased protection, 24 RMs were divided into two groups of animals (n=9 each) and a group of control animals (n=6). Groups 1 and 2 were RhCMV/SIV vaccinated, and Group 3 was left unvaccinated. Vaccination was carried out over 12 months during which RMs were administered the RhCMV/SIV vaccine on weeks 0 and 14. The induction of SIV-specific CD4+ and CD8+ was monitored longitudinally in blood as shown in Figure 3A. The analysis included CD8+ T cell responses to individual MHC-E and MHC-II-restricted 15mer supertopes (Figure 3B).

**Figure 3.**
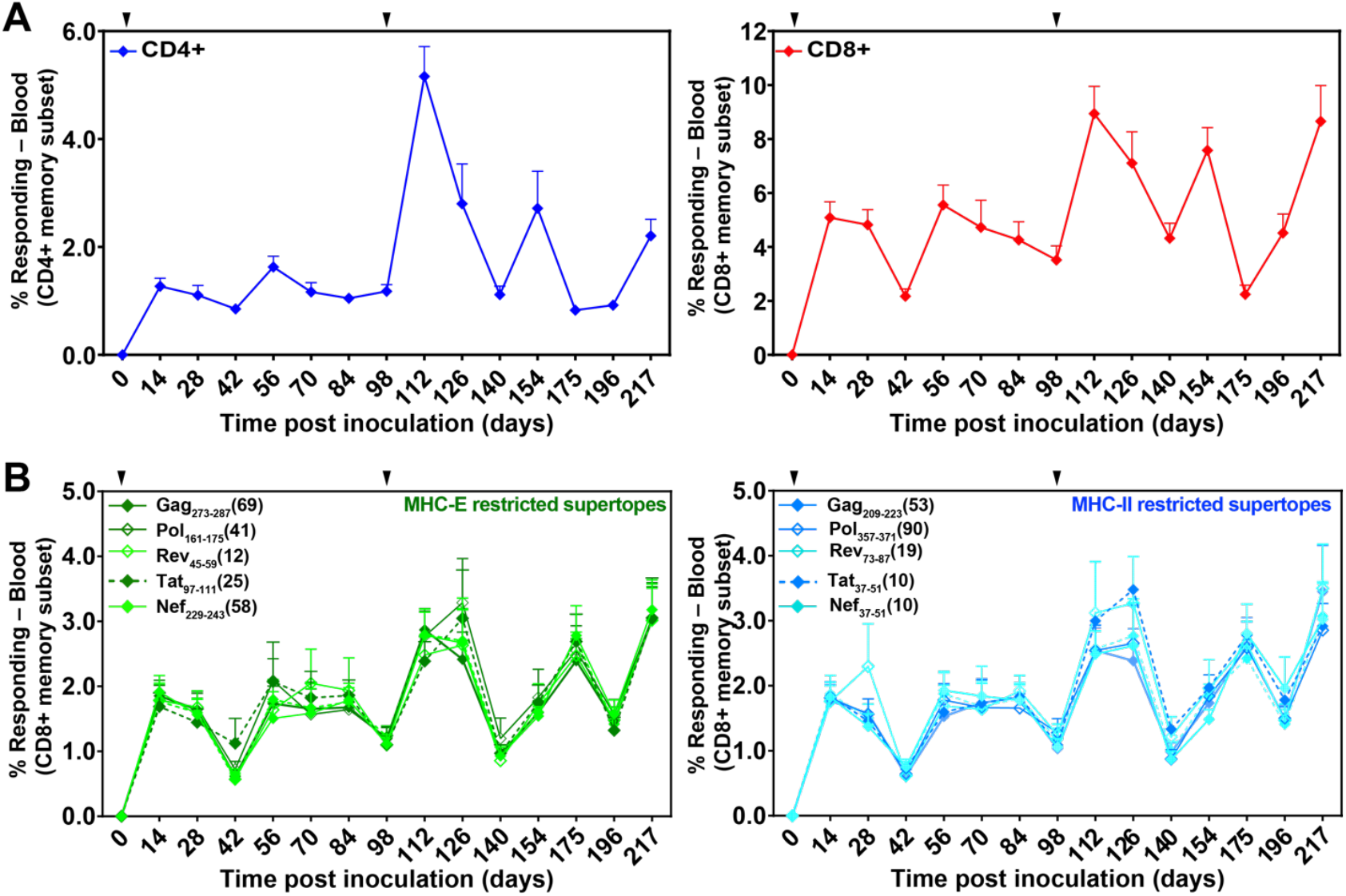
Durability and induction of both SIV-specific T cell responses and unconventionally restricted T cells elicited by 68-1 RhCMV/SIV vaccine vectors. (A) Longitudinal analysis of the overall SIV-specific CD4+ and CD8+ T cell responses in peripheral blood. Responses were determined by ICS analysis (TNF vs. IFN-γ) using whole open reading frame (ORF) mixes of overlapping 15mer peptides (Gag; Rev/Nef/Tat; 5’-Pol) to stimulate PBMCs. The frequency of IFN-γ and/or TNF positive memory T cell responses to each ORF peptide mix were summed to get the overall responses shown in the figure. Vaccinations indicated by the arrowhead above the graph. (B) Longitudinal analyses of CD8+ T cell responses to individual MHC-E-(green; left panel) and MHC-II-(blue; right panel)-restricted 15mer supertopes. Responses were determined by ICS as described in (A).

Twelve months after the first vaccination, Groups 1 and 3 received 3 mg/kg of K11-LS, while Group 2 received 3 mg/kg DEN3, a Dengue virus control antibody, 21–24 days before high-dose (900 FFU) SIVmac239 challenge (Figure 4A, B). Prior to the primary challenge, neutralization ID50s for Groups 1 and 3 were within the desired range of 1:100–1:200 (specifically, geomean values of 1:131 and 1:153, respectively), while Group 2 exhibited no neutralization activity (Figure 4B). After primary challenge, uninfected animals in Groups 1 and 3 were given a second dose of 3 mg/kg K11-LS or control antibody (Group 2) on day 39 and were subsequently challenged again on day 77 (Figure 4A, B).

**Figure 4.**
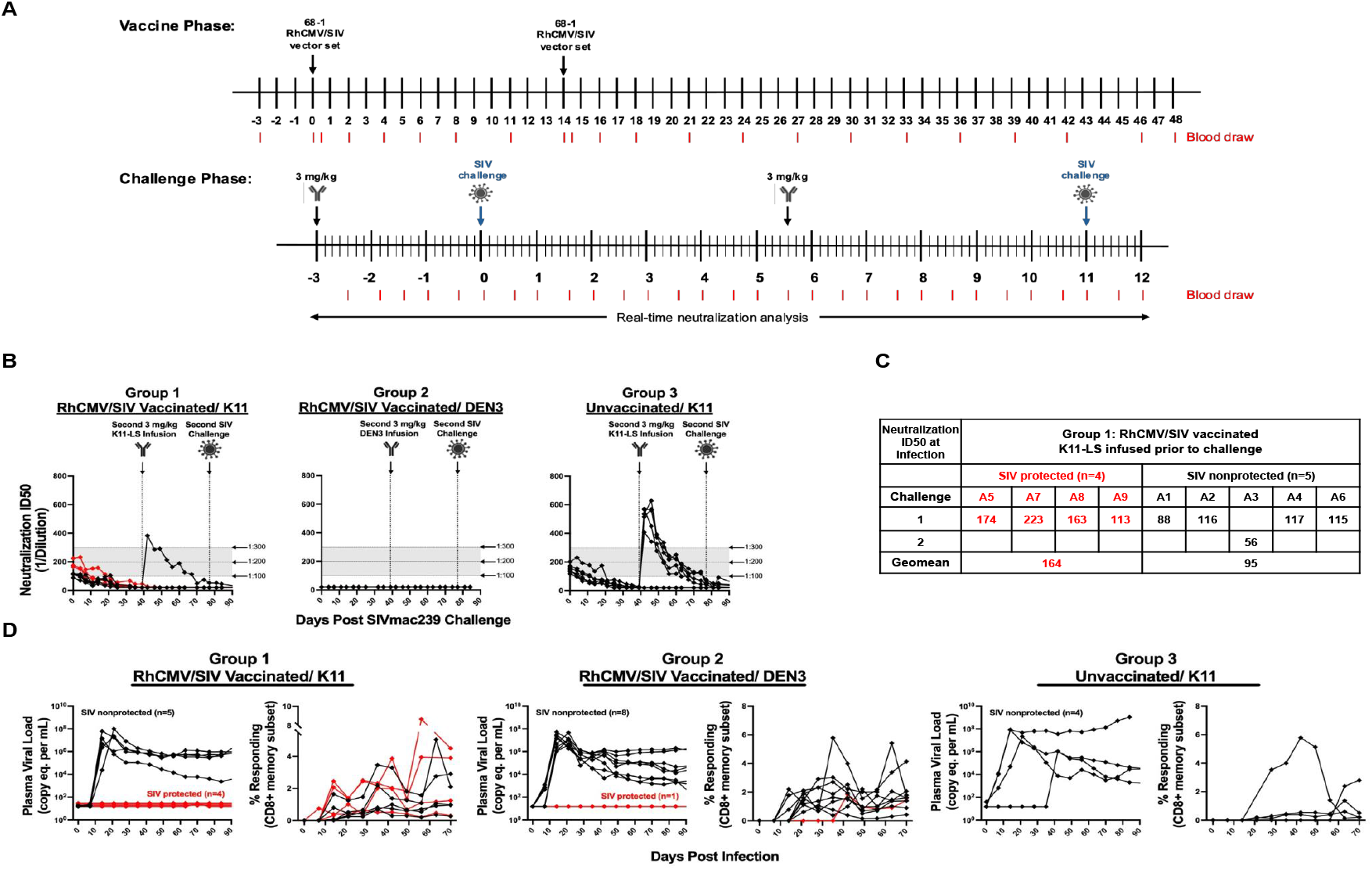
nAb K11-LS facilitates 68-1 RhCMV/SIV vector-mediated protection via replication arrest. (A) The vaccine phase consisted of 12 months during which RMs were administered the RhCMV/SIV vaccine on weeks 0 and 14. During the challenge phase, RMs were administered 3 mg/kg K11-LS (Groups 1 and 3) or control antibody (Group 2) 21 to 24 days before SIVmac239 challenge. Animals A3, B3, B8, C3, C4, C5, and C6 received a second dose of 3 mg/kg K11-LS (Groups 1 and 3) or control antibody (Group 2) 39 days after primary challenge and were challenged a second time on day 77. (B) The neutralization ID50s against SIVmac239 pseudovirus at time of primary challenge were on average 1:130 and 1:160 for Groups 1 and 3, respectively. Group 2 shows no neutralization activity as animals received DEN3 control IgG instead of K11-LS. (C) Within Group 1, nAb titers at time of effective challenge were on average 1:164 and 1:95 for SIV protected and SIV nonprotected RM, respectively. (D) Plasma viral load (PVL) and SIVmac239 Vif-specific CD8 T cells of infected RMs across all three groups. Effective challenge was determined by assessment of plasma virus and *de novo* onset of Vif-specific T cell responses, with infected RMs considered protected if they manifest only Vif-specific T cells (with or without viral blips; red lines) and non-protected if they also manifest sustained plasma viremia (black lines). In Group 3, there are 4 RMs that are non-protected (black lines) and 2 that show no evidence of viral replication and no evidence of viral blips or anti-Vif responses (not shown).

All the RMs in Groups 1 and 2 became infected as indicated by the development of Vif-specific T cell responses following challenge (Figure 4D). In Group 1 (RhCMV/SIV + K11-LS), 4/9 (44%) of the animals showed typical replication arrest as undetectable levels of virus in plasma (Figure 4D). In Group 2 (RhCMV/SIV + DEN3) only 1/9 animals showed replication arrest, the remainder showed robust SIV replication (Figure 4D). Of note, the frequency of replication arrest in Group 2 (11%) was significantly lower than the 50–60% previously observed in studies using repeated limiting-dose viral challenges (Hansen et al., 2011; Hansen et al., 2013a; Hansen et al., 2013b; Hansen et al., 2016; Hansen et al., 2019; Malouli et al., 2021; Verweij et al., 2021). In Group 3, 4/6 animals became infected and showed robust replication of virus; 2/6 animals were not infected and showed no indication of any viral replication as assessed by plasma viral load measurements or anti-Vif T cell responses (Figure 4D). It would appear that these two animals were completely protected by K11, although the serum neutralizing titers were below the 1:300 threshold, or that they had some inherent level of resistance to SIVmac239 infection.

On average, the neutralization ID50 in replication-arrested RMs In Group 1 was 1:164, whereas non-protected RMs had an ID50 of 1:95 at the time of effective challenge when treated with RhCMV/SIV and K11-LS (Figure 4C). This suggests the threshold of neutralization titers necessary for a synergistic efect may be quite high.

## Discussion

We report here a pilot study to investigate potential increased protective activity against SIV infection from combining the orthogonal antiviral properties of RhCMV/SIV vaccine and neutralizing antibody. The number of animals involved was not sufficient to make definitive conclusions, but the study does suggest certain trends that indicate a larger study employing more RMs is merited. With the caveat that it is a single experiment with small animal numbers, this study does suggest that the RhCMV/SIV vaccine may be less effective against a high-dose SIVmac239 challenge than the previously used low-dose viral challenge (Hansen et al., 2011; Hansen et al., 2013a; Hansen et al., 2019; Picker et al., 2023). Thus, replication arrest and protection were observed for only 11% of animals compared with around 60% typically observed, albeit in much larger cohorts of RMs to establish the latter figure. Even with this higher dose challenge, replication arrest and protection were observed in 44% of RhCMV/SIV vaccinated animals when serum neutralizing antibody titers in the range of 100-200 were present as compared to the 11% in the absence of neutralizing antibody. We did observe a higher rate of protection in RMs treated only with neutralizing antibody than expected but the nature of protection was distinct in that it appeared to involve sterilizing immunity with no evidence of virus replication and thus no replication arrest.

Taken together these observations suggest that the suboptimal nAb titers at the time of SIV challenge reduced the size of the initial infection, acting to lower the effective infectious dose. Thus, we would posit that as might be expected from their orthogonal mechanisms of antiviral activity these immune modalities are potentially mutually supportive: RhCMV/SIV vaccination provides cell-mediated replication arrest-type protection when nAb titers are too low to sterilize the challenge, but even a sub-optimal level of neutralization might lower the effective infectious dose enabling vaccine protection when that level is otherwise too high for vaccine-induced MHC-E-restricted CD8+ T cells alone to completely arrest.

What studies should be done next? A first priority would be to provide statistically significant proof-of-concept in RMs, for or against, of the combinatorial efficacy of nAb and RhCMV/SIV-induced MHC-E-restricted CD8+ T cells to protect against SIV. We suggest this could be done in the limiting dose SIVmac239 challenge model by microdosing neutralizing antibody i.e. administering low doses of nAb frequently to RhCMV/SIV-vaccinated animals to maintain subprotective serum concentrations and repeated SIVmac239 challenge. An alternative is to provide an approximately constant level of subprotective nAb by Adeno Associated Virus (AAV) delivery (Johnson et al., 2009; Martinez-Navio et al., 2020) and similarly repeatedly challenging vaccinated animals. Finally, and perhaps most desirably, is a study to identify vaccine constructs capable of eliciting nAbs against SIVmac239, enabling testing of the efficacy of such putative vaccines alone and in combination with RhCMV/SIV vaccination.

## Author Contributions

The study was conceived and designed by J.C., L.J.P., S.G.H., and D.R.B. The experiments were performed by J.C., J.M, R.M.G, S.O., S.F., and D.M.; K.O. and R.B. performed SIV quantitation under the supervision of J.D.L.; AB-A, and R.B. coordinated and implemented RM care, antibody infusions, and procedures. Reagents that were used in this study were produced by J.C., R.B., M.P., K.S. The manuscript was composed by J.C., L.J.P., S.G.H., and D.R.B and all authors reviewed and edited the manuscript.

## Acknowledgements

We would like to express our gratitude to the veterinary staff at OHSU for managing and providing welfare of the rhesus macaques involved in this study. Their expertise and commitment were instrumental in ensuring the health and well-being of the animals, which contributed significantly to the success of this research. Additionally, we are thankful to C. Hughes, A. Selseth, S. Carrizales, R. Sanchez Flores Jr., T. Bennett, C. Pirner, E. Finley, A. Sylwester, N. Hamilton, S. Hagen, and R. Baxter for technical and/or administrative assistance; B. Keele for providing SIVmac239 challenge virus.

